# Thermoregulatory morphodynamics of honeybee clusters

**DOI:** 10.1101/2021.09.13.459268

**Authors:** Jacob M. Peters, Orit Peleg, L. Mahadevan

**Affiliations:** Department of Organismic and Evolutionary Biology, Harvard University; Paulson School of Engineering and Applied Sciences, Harvard University; Department of Computer Science, BioFrontiers Institute, University of Colorado Boulder; Santa Fe Institute, Santa Fe, NM, USA; Department of Physics, Harvard University

## Abstract

During reproductive swarming, honeybees form clusters of more than 10,000 bees that hang from structures in the environment (e.g., tree branches) and are exposed to diurnal variations in ambient temperature for up to one week during the search for a new nesting site. Swarm clusters collectively modulate their morphology in response to these variations (i.e., expanding/contracting in response to heating/cooling) to maintain their internal temperature within a tolerable range and to avoid exhausting their honey stores prematurely. To understand the spatiotemporal aspects of thermoregulatory morphing, we measured the change in size and shape of swarm clusters over time and the internal temperature profiles in response to dynamic temperature ramp perturbations. We found that swarm clusters can achieve a twofold increase/decrease their volume/density when heated from 15°C to 30°C, but they do not reach an equilibrium size or shape when held at 30°C for 5 hours, long after the core temperature of the cluster has stabilized. Furthermore, the changes in cluster shape and size are hysteretic, contracting in response to cooling faster than expanding in response to heating. Although the contact diameter of the cluster increased continuously when the swarm is heated, the change in length of the swarm (base to tip) over time is non-monotonic. Consequently, the aspect ratio of the swarm fluctuated continuously even when held at a constant temperature. Taken together, our results quantify the hysteretic and anisotropic morphological responses of swarm clusters to ambient temperature variations while suggesting that both mechanical constraints and heat transfer govern the thermoregulatory morphing dynamics of swarm clusters.

## Introduction

Many social hymenopterans (i.e., ants, wasps and bees) engage in a class of collective behaviors in which individuals link their bodies together to form self-assembled structures such as rafts [13], bridges [19], droplets [2] and clusters [8]. These behaviors often serve adaptive functions, aiding in flood survival, predator evasion, food transport, navigation of complex environments, etc. (see Anderson et al. [1] for a review). The microenvironment within these aggregates is an extended phenotype of the colony that can give rise to emergent physiological functions. For example, bivouacs and clusters formed by army ants [6] and honeybees [9] are able to maintain relatively stable internal temperatures despite large variations in ambient temperature. Studying these behaviors can provide insight into how the interplay between collective behavior and microenvironment can lead to colony-level physiology.

In honeybee swarm clusters that can have of the order of ≈ 10, 000 bees that settle on a tree branch (Fig. 1), exposure to a variable environment for days while scouts seek new nest sites [21] causes the cluster to modulates its morphology in an attempt to maintain a relatively stable microenvironment. For example, when exposed to rain, the bees on the surface of the cluster adjust their posture to collectively shed water much like a tiled roof does [4], while in response to mechanical perturbations (such as being shaken by the wind), the cluster collectively flattens out and increases its attachment area to reduce mechanical strain on the cluster and increase its overall stability [18]. Swarm clusters also modulate their morphology in response to changes in ambient temperatures in order to maintain relatively stable internal temperatures, expanding to dissipate heat when ambient temperatures rise and contracting to conserve heat when ambient temperatures drop [8, 9]. Here, we investigate this behavior by quantifying the extent of thermoregulatory adaptation via morphological variation.

**Figure 1:**
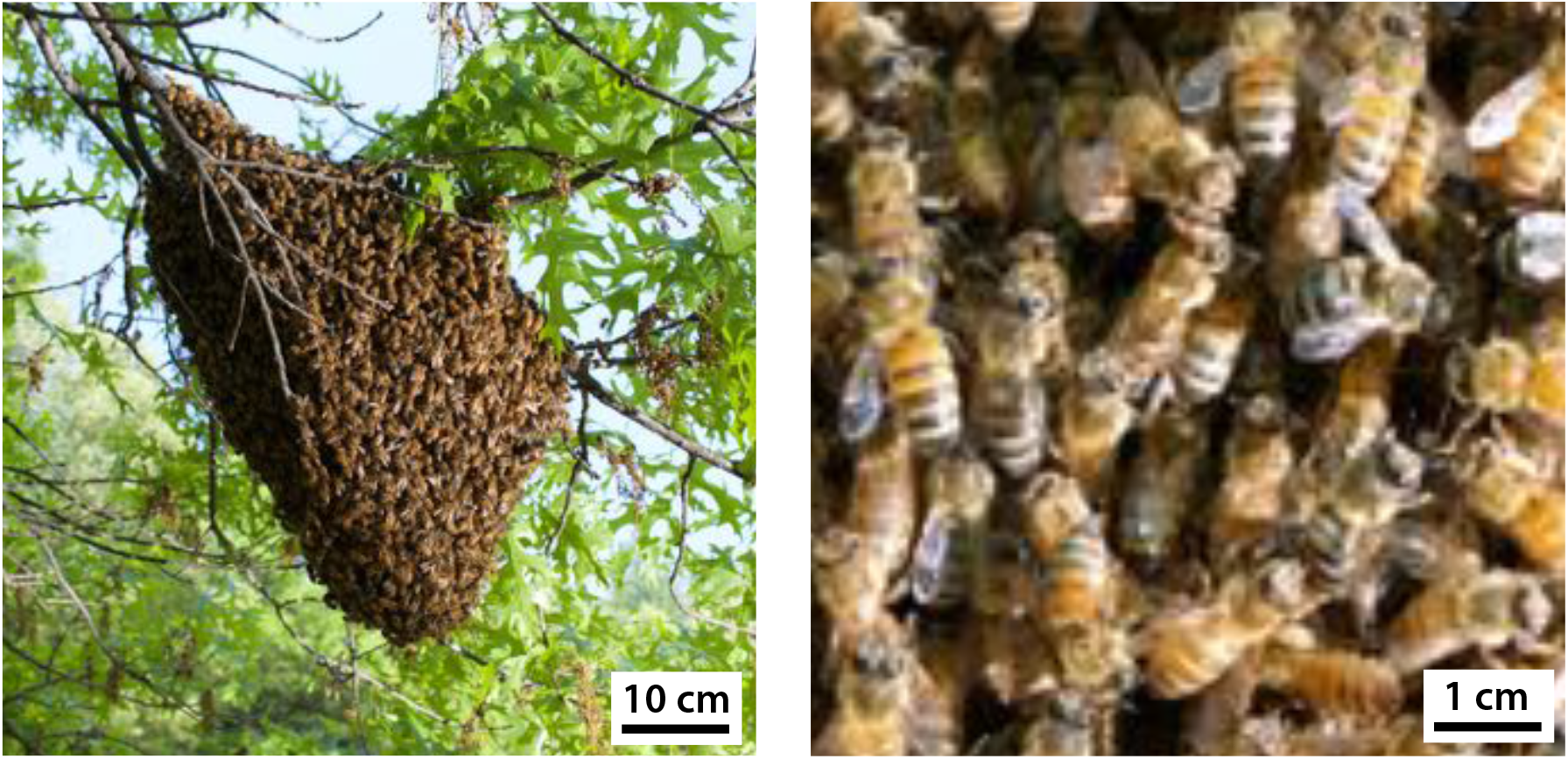
A feral honeybee swarm at the Concord Field Station (left). Close-up view of bees on the surface of the cluster (right).

Early experiments pioneered by Heinrich [8] measured the core and surface temperature swarm clusters across a broad range of fixed ambient temperatures (1 − 35°C). In large clusters(> 15000 bees), the core temperature of large swarms was relatively stable across this range at 35 − 36°C, while in small swarms (*<* 10000 bees) the temperature was more variable, ranging from 20 − 35°C. When the ambient temperature was below 17°C, the surface temperatures were actively maintained at ≈ 17°C through shivering (i.e., vibration of the thorax) in order to prevent chill coma (i.e., cold-induced immobilization). At ambient temperatures greater than ≈ 17°C the surface temperatures were maintained a few degrees above ambient temperature. Heinrich’s observations showed that the cluster increased its volume and surface area dramatically as ambient temperatures rose from 17 to 26°C, allowing more metabolic heat to dissipate and avoiding overheating. As ambient temperature dropped below 17°C the cluster reduced its surface area and increase its density to conserve metabolic heat. The change in size/shape below ≈ 17°C was subtle suggesting that bees may approach their maximum density. Heinrich also measured the collective metabolic rate of the cluster and showed that oxygen consumption increased linearly as ambient temperature rose above 17°C and increased even more steeply as ambient temperatures dropped below 17°C. Taken together, these results suggested that the swarm modulates its metabolic rate below 17°C to prevent chill-coma and modulates its morphology above 17°C to minimize energetic costs and to prevent overheating [8].

Thermoregulation in swarm clusters is an intriguing multi-scale control problem wherein individual bees (2cm) presumably sense/respond only to local information and yet collectively they achieve relatively stable cluster (10 − 30cm) temperatures. A simple way to explain this global regulation is to recognize that it can emerge from coupling between local behaviors, the (geometrical and topological) structure of the cluster and the environmental physics of heat transfer that both enables and constrains function, as studied quantitatively in bees, termites etc. [15, 10, 3]. The specific problem of thermoregulation in bee clusters has been the subject of many theoretical studies [16, 17, 11, 5, 14, 23, 22, 15]. All models involve the conduction and convection of heat in a metabolically active materials and some include “thermotactic” behaviors in which bees modulate their activity according to local temperature [14, 23, 22, 15]. These models focus primarily on the steady state behavior of the loaded network of bees that are assumed to be able to rapidly equilibrate with the environment by instantaneously breaking and reforming attachments between neighboring bees. A natural question that remains open is that of the morphodynamics of thermoregulation.

To understand the dynamics of swarm thermoregulation, here we focus on measuring the change in size, shape and temperature profiles of swarm clusters over time in response to dynamic temperature ramp perturbations. This allows us to measure the dynamical morphological response of the system and thus shed light on the temporally transient and spatially anisotropic collective behaviors that arise from local responses of individual bees.

## 1 Methods

### Artificial swarms

The artificial swarm clusters used in experiments consisted of packages of honeybees purchased from New England Beekeeping in Tyngsboro, Massachusetts in the spring of 2017 and 2018. These packages consisted of approximately 1 kg of European honeybees *Apis mellifera* and a caged queen. For at least two days prior to the experiments the bees were fed *ad libitum* 2:1 sugar water solution, which is known to induce natural swarming behavior [20].

### Experimental Apparatus

The experimental setup consisted of a 1.2 × 1.5 × 1.2 m box constructed from 3.8 × 8.9 cm dimensional lumber and insulated with a double layer of press-fit 2.5-cm Styrofoam sheets (Fig. 2A). Two window air conditioners were inserted into holes at the bottom left and right corners of one of the walls of the box, and three small space heaters were placed in the corners of the box. Four temperature sensors were suspended in the airspace of the box and connected to an Arduino microcontroller. The heaters and air conditioners were toggled on and off in response to this temperature feedback in order to maintain a desired set-point temperature. This set-point temperature was modulated by a Matlab program to achieve controlled heating and cooling cycles.

**Figure 2:**
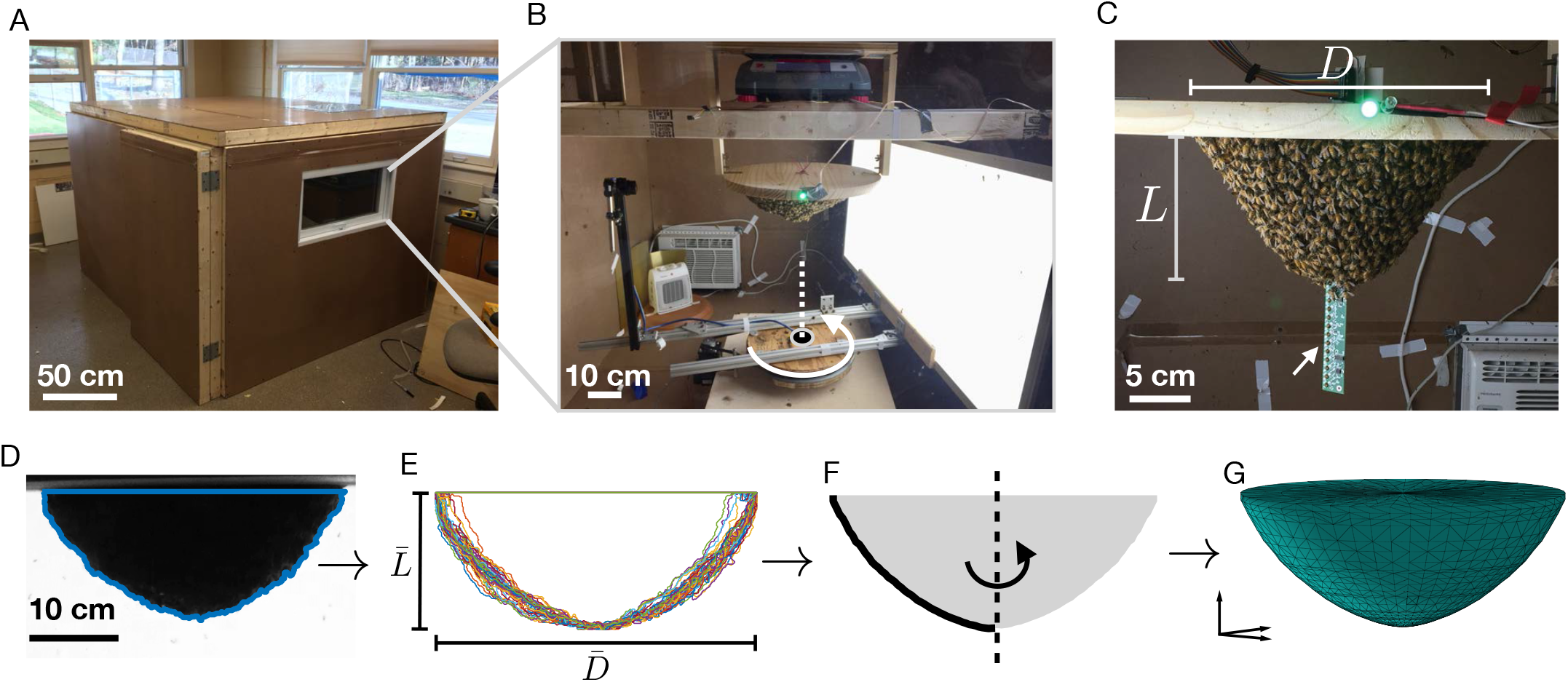
Experimental setup for quantifying the thermoregulatory response of swarm clusters. (A) Swarms were housed in a large insulated box, outfitted with heaters and coolers that were toggled on and off to produce controlled temperature perturbations. (B) Within the box, a board (to which the swarm was attached) was suspended from a scale. A motor was used to revolve a camera and an opposing back light 360° around the swarm. (C) In some trials, a PCB with a 32-sensor array (white arrow) was inserted in the swarm cluster to record the temperature gradient within the swarm. Bars corresponding to *D* and *L* indicate the diameter and length of the cluster, respectively. (D) During each rotation, 40 images of the swarm were captured from different angles. (E) The swarm was segmented from the background of each image using thresholding. (F) The silhouettes of the 40 images were transformed so that diameter *D* and length *L* matched the average values for the whole set, 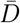 and 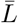. (G) An average shape was calculated from the 40 silhouettes, and this shape was revolved around the *z*-axis to obtain an axisymmetric, 3D approximation of the swarm’s shape.

Two wooden struts were used to support a scale (Ohaus Ranger 3000) approximately 10 cm from the roof of the box, and a 60-cm diameter wooden board was hung from the scale (Fig. 2B). The queen cage was fixed to the underside of this board with wires. The rest of the bees were allowed to form a cluster around the queen until the mass reported by the scale was stable, indicating that all of the bees had joined the swarm cluster. The bees were allowed to fly freely, both within the box and to fly in/out of a 0.25 × 1 m window in the side of the box which connected to the outdoors. This allowed the swarm to send scouts in search of new nest sites. In preliminary attempts using a completely closed box we found that the bees would continuously search for an opening in the box to escape from if they were not given freedom to fly in/out of the window and the mass of the cluster would not stabilize.

Each experiment was 10-16 hours long. Initially, the air temperature in the box was held at 15°C for 4 hours, then ramped up to 30°C for 5 hours, and finally ramped back down to 15°C for an additional 4 hours. Three heating/cooling rates were tested: 33.4°C/hr (*n* = 6), (B) 8.6°C/hr (*n* = 9) and (C) 4.3°C/hr (*n* = 5). On warm, sunny days the bees would occasionally fly off as if to a new nest site only to return to cluster around the caged queen. This behavior became so disruptive that some of the trials were conducted over night. See Table S1 for a list of dates/times of individual trials as well as for the ID of the swarms used.

### Image acquisition and analysis

A machine vision camera (1.3 MP, Point Grey Chameleon3) was placed 50 cm from the board supporting the cluster and an LED back light (50 × 100 cm) was placed orthogonal to the camera on the other side of the swarm. During each data collection bout, the back light was switched on and the camera was rotated 360 degrees around the swarm (Fig. 2B, Movie S1). Images were captured every 9 degrees of rotation resulting in 40 unique views of the swarm (Movies S1 and S2). When the revolution was complete, the back light was turned off to avoid luring bees away from the cluster. This filming routine was executed approximately every 4 minutes during the experiment. The mass of the swarm was also continuously recorded throughout the experiment.

Prior to analysis, each image of the swarm was undistorted using Matlab’s checkerboard calibration tool (to correct for wide-angle lens geometry). The checkerboard was placed orthogonal to the camera at the axis of rotation so that images in that plane could be converted from pixels to centimeters. A custom Matlab image segmentation program was then used to define a wire frame silhouette of the cluster in each of the 40 views (Fig. 2D). Because the cluster was rarely centered at the camera’s axis of rotation, the silhouette of the cluster appeared to loom and recede during the camera’s rotation. To correct for this, each silhouette was transformed such that its dimensions (i.e., diameter and length) equaled the average dimensions of the set of 40 silhouettes (Fig. 2E). The silhouettes were then averaged to achieve a single 2D approximation of the profile shape of the cluster centered at the axis of rotation (Fig. 2F). The diameter *D* of the swarm and the length *L* cluster were determined using this averaged silhouette. The perimeter of the averaged silhouette was revolved around the vertical axis to create an axisymmetric, 3D approximation of the shape of the cluster, and to determine its surface area *A* and volume *V* (Fig. 2G, Movie S4). We chose to use this method rather than hull reconstruction because it was more robust to bees flying in front of the camera (which was a persistent problem).

### Recording internal temperature profiles

We used a custom sensor array (Fig. 2C) to record the internal temperature profile of a single swarm cluster over the course of two experiments with two different heating/cooling rates (i.e., 33.4°C/hr and 8.6°C/hr). We also attempted to record the internal temperature of a cluster at a heating/cooling rate of 4.3°C/hr, but we later noticed that the bees had begun building comb on the sensor array. Bees elevate their body temperatures in order to build comb, which would confound our experiments. Therefore, we excluded this trial.

The sensor array consists of a printed circuit board (PCB) with 32 temperature sensors (i.e., 10k-ohm glass bead NTC linear response thermistors) spaced every 0.89 cm along the length of the PCB. To ensure that the sensors measured the temperature of the air and/or bees within the cluster rather than the surface temperature of the PCB itself, the sensors were suspended by their lead wires inside of 0.32 cm holes in the PCB, such that the glass bead was not in direct contact with the PCB. In order to avoid self-heating, a digital switch was used to ensure that the sensors received power only while the measurement was being made (i.e., about 500ms). Measurements from all 32 sensors were recorded every 10 seconds throughout the experiment. The sensors were calibrated by running a temperature ramp experiment without the swarm present and fitting a linear model relating the output voltages to air temperature.

## Results

The scaled change in each response variable (i.e., mass 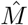, volume 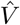, density 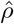, area *Â*, diameter 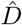, length 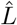) is is plotted in Figure 3A-C. Each variable *y* is scaled by dividing the change in that variable by the initial value, *ŷ* = (*y*(*t*) − *y*(0))/*y*(0). This removes the dependence on the sizes of the clusters, which were variable. See Table S1 for initial conditions for each response variable.

**Figure 3:**
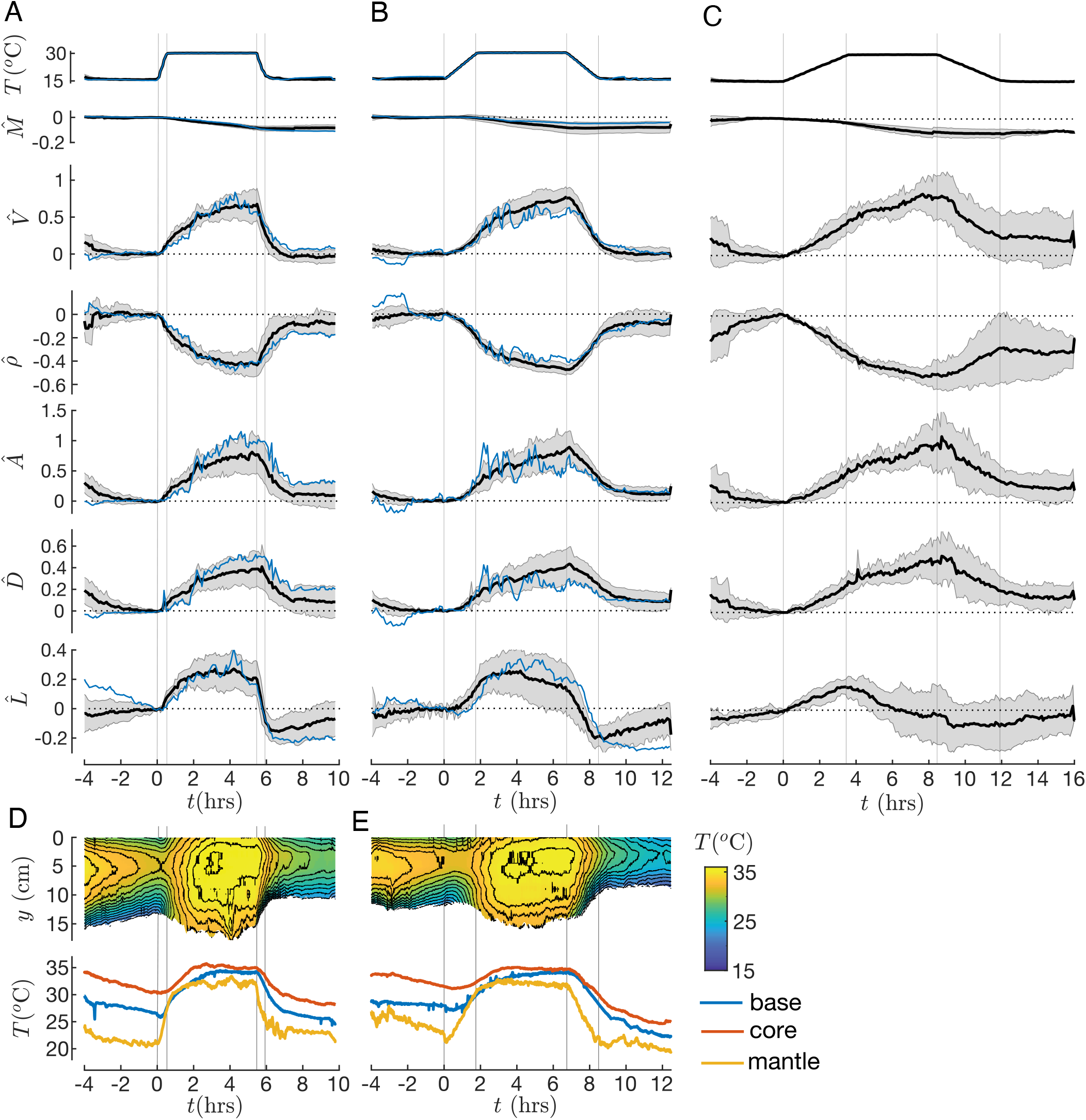
The dynamic response of honeybee swarm clusters to controlled temperature ramp perturbations. Each response variable *y* is scaled such that *ŷ* = (*y*(*t*) − *y*(0))*/y*(0). The scaled change in volume 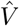, surface area *Â*, diameter 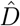, length 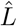, density *rĥo* and mass 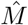 are plotted for 3 treatments with varying heating/cooling rates: (A) 33.4°C/hr (*n* = 9), (B) 8.6°C/hr (*n* = 9) and (C) 4.3°C/hr (*n* = 5). The treatment mean is plotted in black and the gray error band indicates standard deviation. In A and B, the blue line indicates the response for trials in which a sensor array was used to record internal temperature data (as shown in D-E). Continuous measurements of the temperature profile within swarm clusters. Internal temperature data is provided for one trial for two heating/cooling rates 33.4°C/hr (D) and 8.6°C/hr (E). The heatmap (top panel) represents the temperature at various positions *y* within the cluster over time *t*, where *y* = 0 is base of the swarm and *y* increases towards the tip of the swarm. Data recorded from sensors external to the swarm are omitted. In the lower panel, the temperature at the base, core and mantle of the swarm are plotted over time.

### Changes in the Mass of Cluster

The clusters lose 8-10% of their initial mass over the course of a heating/cooling experiment (Fig. 3A-C). Although we did not quantify the source of this weight loss, we noticed several potential sources of weight loss. First, we noticed that wax scales accumulated on the floor of the box during the experiments. Honeybees regularly produce wax scales in preparation for colonizing a new nest site [7]. In addition, some of the loss may be attributed to attrition of bees, as some dead bees accumulated on the floor of the box. However, this did not appear to be enough to account for the 8-10% weight loss. It is also possible that some of the weight loss resulted from bees leaving the box and failing to return to the swarm, though scouts appeared to move in and out of the box with ease. In fact, on a few occasions between experiments the entire swarm flew out the window (likely toward a newly selected nest sight), only to return to the box after discovering that their queen was not with them. In these instances, the mass of the swarm was similar before and after its departure, suggesting that the bees did not struggle to find their way back to the box.

Interestingly, the rate of weight loss was affected by the ambient air temperature (Fig. 3A-C, panel 2). The clusters lost weight at an average rate of 2.8%/hr during the 30°C plateau and was virtually constant during the final 15°C plateau. This temperature dependent weight loss is not unexpected, as [9] demonstrated that the metabolic rate of swarms increases two fold from 15°C to 30°C. Both behavioral mechanisms of weight loss (e.g., bees getting lost) and physiological mechanisms of weight loss (e.g., wax excretion, metabolic water loss, respiration, death of individuals) are likely to be affected by metabolic rate.

### Morphological Response

To characterize the morphology of the swarm cluster, we measured the volume and area of the cluster as well as its diameter and height as a function of the imposed temperature perturbations, as shown in Figure 3A-C (see Table S2 for summary statistics). When the temperature was raised from 15°C to 30°C the clusters nearly doubled in volume, but never actually reached a steady state, even after a 5-hour heating phase. However, when the temperature was dropped to 15°C during the cooling phase, the volume *V* returned to the initial volume *V*_0_ within 2 hours for treatments 1 and 2, but did not quite return to *V*_0_ for treatment 3. Similar dynamics were observed for surface area, although it did not quite return to its initial value by the end of the experiments. As expected given the doubling in cluster volume, the density of the clusters was approximately halved by the end of the heating phase.

We expected both the diameter and the length of the cluster to increase monotonically during heating and decrease monotonically during cooling. However, while diameter increased monotonically during heating and reached nearly 50% of its initial value, length showed a non-monotonic response. In fact, length initially increased upon heating but reached a peak and began to fall before the end of the 30°C plateau. During cooling, diameter decreased monotonically and nearly reached its initial value by the end for the experiment. Length, however, rapidly decreases upon cooling, overshooting its initial value and finally begins to rise again during the 15°C plateau, almost reaching its initial value.

### Internal Temperature Variation

Since the morphological response is correlated with the internal temperature profile of the cluster, we measured this along the central axis of a swarm cluster during experiments at two different heating/cooling rates (i.e., 33.4°C/hr and 8.6°C/hr), using the setup is depicted in Figure 3F, and presented in Figure 3D-E. In both experiments, the core of the swarm was generally less variable and warmer (30-36°C) than the mantle (19-33°C) which typically remained several degrees above the ambient temperature when above 15°C. These results are in agreement with previous reports from Heinrich [9] and are consistent with the hypothesis that the bees attempt to maintain temperatures between a lower bound (≈ 17°C) and an upper bound (36°C).

At the beginning of a heating protocol, the mantle began to warm immediately, but there was a lag in the onset of heating at the core of the swarm of about 0.5 hrs for the 33.4°C/hr experiment and 1 hr for the 8.6°C/hr. This lag was reflected by a corresponding lag in the morphological response of the cluster as evidenced by the delayed increase in volume and surface area. These observations are consistent with the morphological response to heating being triggered/driven by bees in the interior of the swarm responding to a gradually rising local temperature due to the conduction, diffusion and convection of heat from the boundary.

However, the core temperature of the swarm was significantly lower at the end of that experiments than the initial core temperature. This suggests that the swarm cluster is particularly susceptible to cooling after extended periods at high temperatures, consistent with the fact that clusters reduced their density by about 50% and increased their aspect ratio during the 30° plateau. The cluster is able to rapidly decrease its length in response to cooling in order to increase its density and conserve heat. However this results in a sub-optimal, high aspect-ratio shape (short and wide) with a high surface area-to-volume ratio. When the cluster is already at high density, recovering a lower aspect ratio shape with a low surface area-to-volume ratio seems to be a slow process. The slow return to a compact shape may be responsible for the observed over-cooling of the core.

### Hysteresis

To better understand the nonlinear response of the cluster to dynamic temperature perturbations, we fit the mean 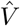 and *Â* to simple first-order kinetic response functions associated with the slowest time scale *τ* in the relaxation processes at play, assumed to be of the form *dr/dt* = (*s*(*t*) − *r*(*t*))*/τ*, where *r*(*t*) is the response to a given stimulus *s*(*t*) (see Fig. 4A-B).

**Figure 4:**
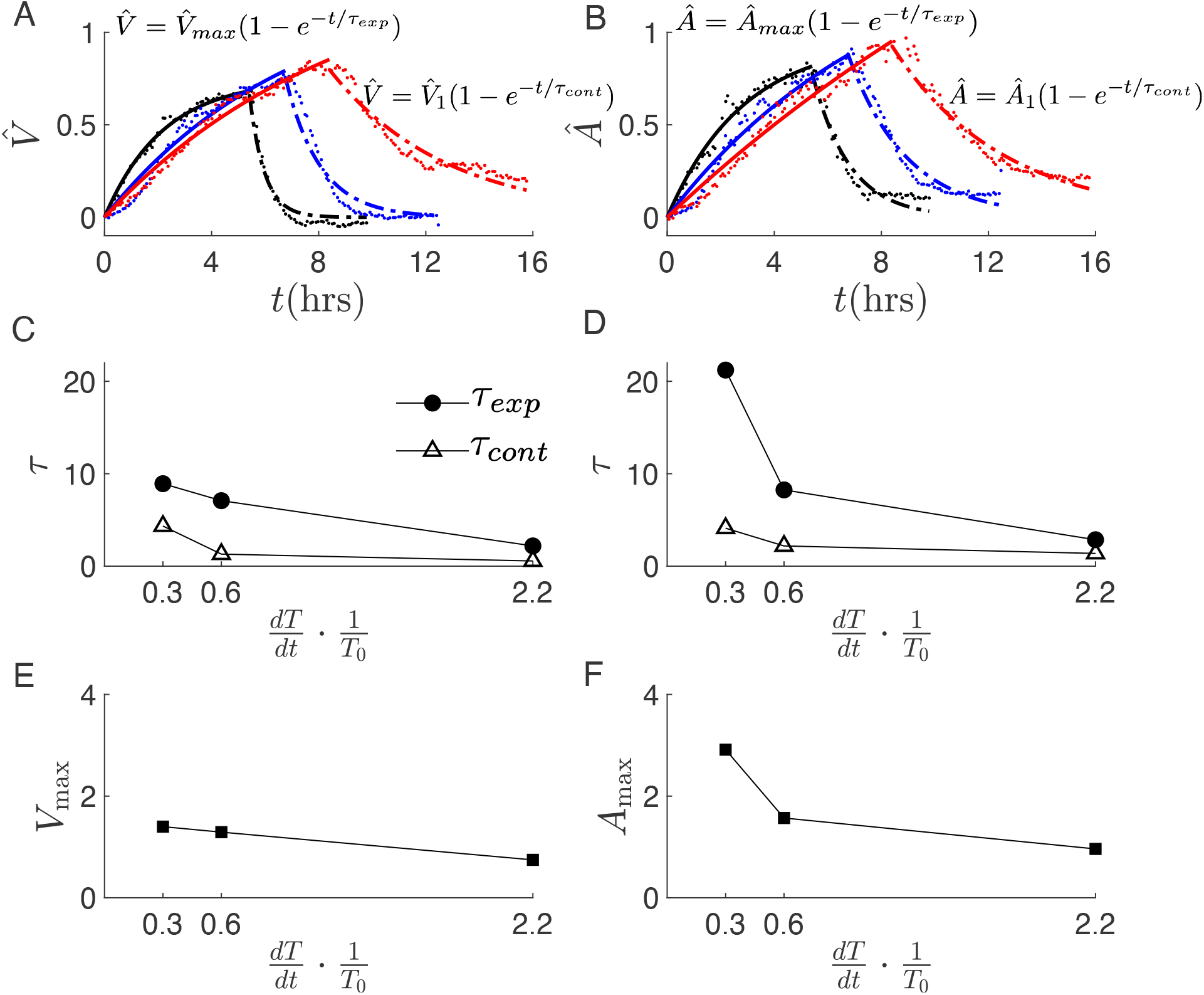
Hysteresis in the thermoregulatory expansion (solid line) and contraction (dotted line) responses of swarm clusters. The mean (relative) change in cluster (A) volume 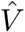 and (B) area *Â* over time for each treatment group: 33.4°C/hr (black), 8.6°C/hr (blue) and 4.3°C/hr (red). The rising edge of the response (expansion) is fit to 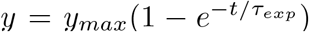 and the falling edge of the response (contraction) 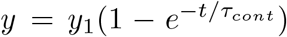. The timescale for the rising *τ*_*exp*_ (solid circles) and falling *τ*_*cont*_ edge (open triangles) are plotted across heating/cooling rates, which are represented by the dimensionless parameter 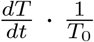. Large values of *τ* represent slow responses and small values indicate faster responses.

For the response to an increase in the temperature (i.e., expansion of the cluster), we fit the response to

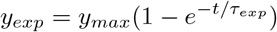

where *y*_*exp*_ is the response variable, *τ*_*exp*_ is the time constant of the response, and *y*_max_ is the value at which *y* would eventually saturate if allowed enough time. For the response to a decrease in the temperature (i.e., contraction of the cluster), we fit the response variable to

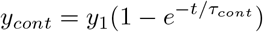

where *y*_1_ is the value of y at the start of the cooling ramp. For all trials, the timescale of volumetric expansion was higher than that of contraction, indicating that expansion occurs more slowly than does contraction (Fig. 4C-D). This difference in expansion/contraction timescales was more pronounced for slow heating/cooling treatments than for fast heating/cooling treatments. We observed an even more dramatic difference in surface area expansion and contraction timescales. Taken together, this suggests that the expansion/contraction response shows strong hysteresis.

Neither 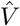 nor *Â* reached saturation during the five hour long 30°C plateau. The parameters 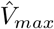 and *Â*_*max*_ from the response model are estimates of the maximum values that 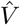 and *Â* would have reached if the 30°C plateau was sufficiently long (Fig. 4E-F). We expected that *V*_*max*_ and *A*_*max*_ would be constants, however they appear to decrease with the heating rate of the treatment. This suggests that the final volume and surface area of the cluster are greater if you heat them more slowly.

### Anisotropy and Shape Changes

The changes in diameter 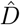 and length 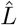 are plotted together in Figure 5A-C to highlight their different but coupled dynamics. The scaled change in aspect ratio 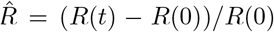 is plotted in Panel 2, where aspect ratio is the ratio of width to length *R* = *D/L*. Key transitions in the shape/size of the clusters are denoted in Figure 5 by roman numerals (I-V). We note that these transitions are related to the nature of the heating/cooling ramps, but they are not necessarily coincident. The aspect ratio stays approximately constant at the start of each experiment as diameter and length increase together (I-II). However, length begins to decrease before the end of the 30°C plateau causing the aspect ratio to increase (II-III). At the onset of the cooling ramp, aspect ratio undergoes a sharp increase as cluster length decreases dramatically faster than diameter leading to an even flatter shape (III-IV). This sharp increase is most pronounced for the fastest treatment and is nearly absent for the slowest treatment. Finally, the cluster begins to slowly return to its original shape during the 15°C plateau (IV-V). Aspect ratio does not quite return to its initial value by the end of the experiment (V).

**Figure 5:**
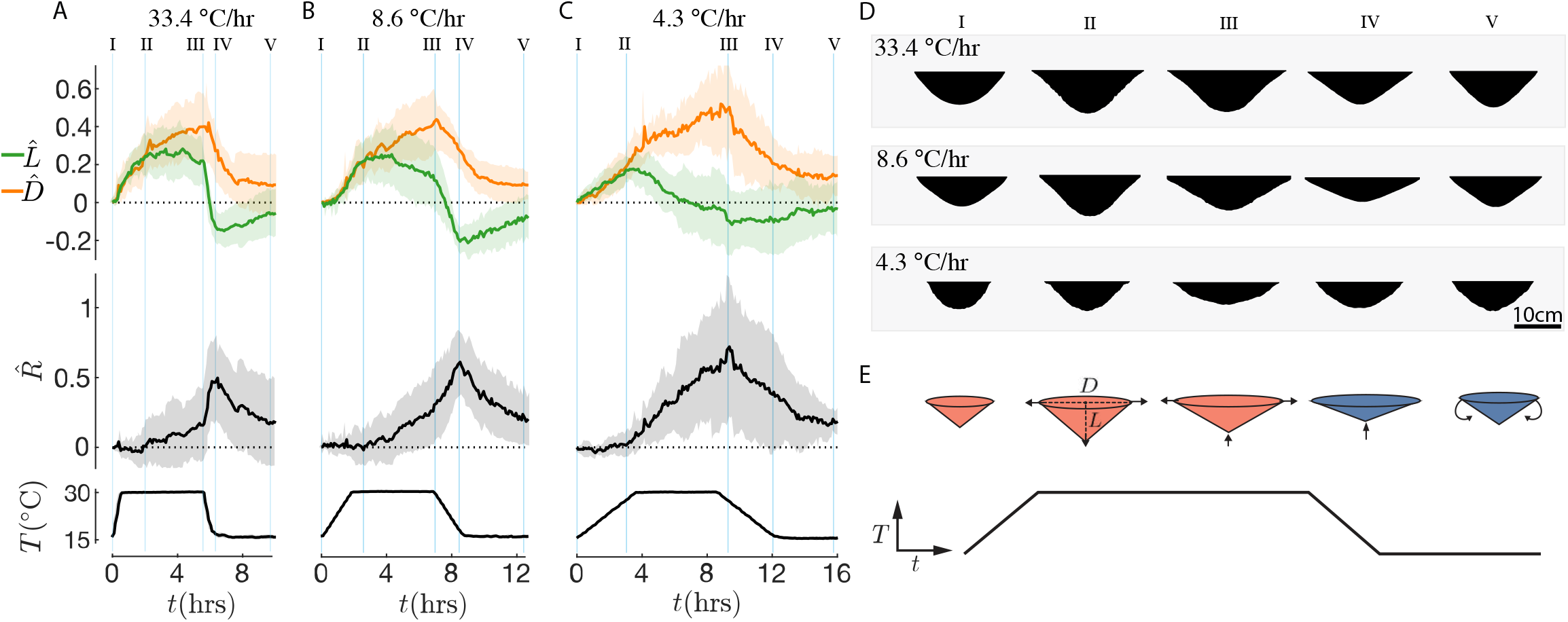
Dynamic changes to the shape of swarm clusters in response to temperature perturbations. In the top panel, the change in length 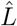 (green) and 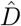 (orange) are plotted over time for each treatment group (A-C). Each line represents the mean and error bands spans the mean *±* standard deviation. In the second panel, The change in the aspect ratio 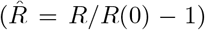 relative to the initial aspect ratio (*R* = *D/L*). The temperature perturbations are plotted in the bottom panel to for reference. Key transitions are denoted by roman numerals (I-V). They are not necessarily coincident with the transitions in the heating/cooling ramps. (D) Averaged silhouettes of clusters at these transitions are are shown for a representative trial from each treatment group. (E) The shape of the swarm is represented with a series of cones which change dimensions over time in response to a temperature ramp perturbation (depicted below). In response to heating (red cones), the swarm initially expands in diameter and length. However as diameter continues to increase, length starts to decrease, likely to prevent the density from reaching a minimum density below which the swarm would break. During cooling (blue cones), the length of the cluster rapidly shortens. The unloaded bees at the surface likely move upward to fill gaps in the swarm. Initially bees at the edge of the clusters base are not able to move toward the center of the cluster because they are loaded (or supporting the weight of other bees). But as the bees below them move inward, they become free to shift as well. This may be why the decrease in diameter lags behind the decrease in length.

These dynamics suggest that there are two distinct phases of the response to heating (Fig. 5E). In the first phase, as heating begins the bees begin to increase their volume and thus decrease their density relatively quickly, allowing for convective cooling keeping the shape of the cluster invariant. This of course cannot continue forever owing to the mechanical constraints of keeping the cluster contiguous and strong enough to support itself. The second phase involves changing the shape of the cluster subject to keeping the volume (and the mass) constant, but occurs much more slowly.

To quantify the morphological changes, we recall that for a generalized conically shaped cluster with a base of radius *r* and height *H*, we can write

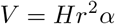

where *α* is a constant associated with the shape of the cluster (for a right cone *α* = *π/*3, while for a cylinder *α* = *π*). The diameter and length of the cluster initially increase together to achieve this increase (decrease) in volume (density), preserving the shape but increasing the size. However, as the density of the cluster approaches a minimum (below which the structural integrity of the swarm would likely be compromised), diameter and length can no longer increase together. Instead, the cluster increases in diameter and decreases in length, i.e. the shape now changes. This allows the surface area to continue to increase while slowing the increase (decrease) in volume (density), thereby preventing mechanical failure. The surface area of the cluster is given by

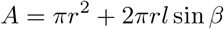

where *β* is the half-angle of the cone. An additional benefit of this change in the shape (i.e., flattening of the cluster) is the advantage that it proffers from a mechanical perspective, consistent with a recent experimental and theoretical study [18]. This study demonstrated that flatter swarm clusters with broad attachment areas are more stable than elongated clusters with small attachment areas (Fig. 6a) because they more efficiently distribute the load and further that bees are able to increase their aspect ratio (and attachment area) in response to imposed mechanical perturbation. These responses of the bee cluster shape to temperature and mechanical loads suggest that in general the shape is a consequence of the ability of individual bees to respond using fast limb extension or retraction movements and the slower body repositioning based both on local temperature and mechanical information.

**Figure 6:**
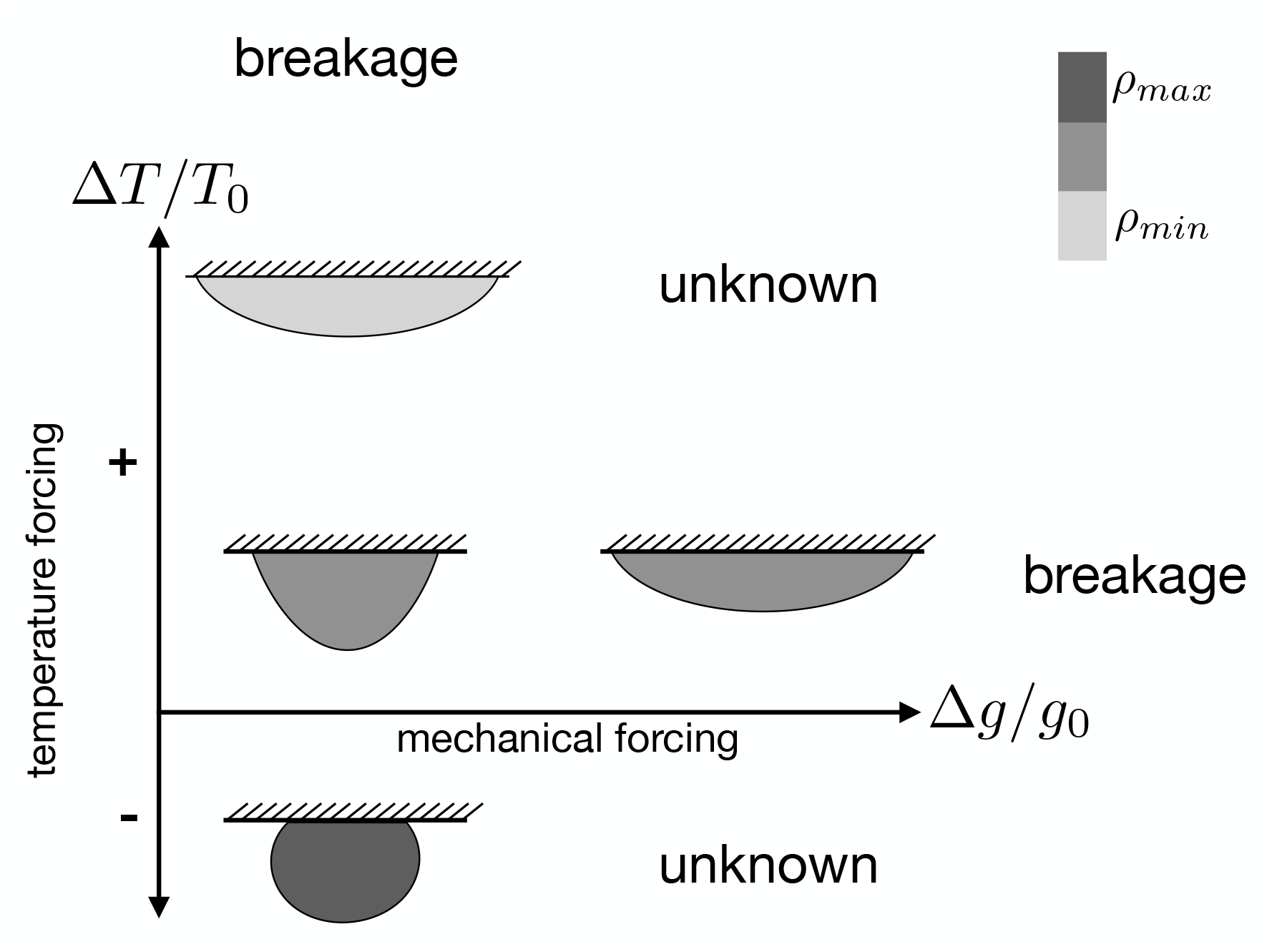
A schematic of the known phase space of honeybee swarm cluster density/shape according to this and previous studies [18, 9]. Mechanical forcing on the x-axis is reported as Δ*g/g*_0_ where *g*_0_ = 9.8m*/*s2, the acceleration due to gravity. Temperature forcing on the y-axis is reported as Δ*T/T*_0_ where *T*_0_ = 17°C. Below *T*_0_ the cluster reaches a maximum density and thermoregulates primarily by modulating metabolic heat production through shivering. The shape of the cluster is represented by cartoons (not to scale) and the density of the cluster is represented by the color axis. How the swarm integrates both temperature information and mechanical information to maintain stability when temperature and mechanical forcing are decoupled is unknown. This provides a unique opportunity for future study of multi-sensory integration at the colony scale.

## Discussion

Our study complements and build on the pioneering experimental work of Heinrich [9] on honeybee swarm clusters that demonstrated their ability to maintain a relatively stable internal temperature despite dramatic fluctuations in ambient temperature. The proposed mechanism for this was postulated to be the modulation of metabolic heating (through shivering) at low temperatures and by controlled heat dissipation at high temperatures (through collective expansion and contraction of the cluster). This inspired many mathematical models which treat honeybee swarm clusters as a metabolically active material which generates heat and also responds by changing its size, shape and density [16, 17, 11, 5, 14, 23, 22, 15] in response to local temperature information. These models demonstrate that thermoregulation is possible even without direct communication among bees due to physical coupling between individuals. However, these studies typically consider the steady-state of the swarm because previous experiments measured the temperature, metabolic activity and size of the clusters at fixed temperatures [9, 8].

Our work generalizes this and contributes to the understanding of collective thermoregulation in swarm clusters by measuring their morphological responses to time-varying temperature perturbations. Our results show that expansion and contraction of swarm clusters has two phases, a fast and a slow phase (Fig. 5).

At the start of the heating ramp the clusters initially increase rapidly in both diameter and length (phase 1), but then the cluster begins to shorten in length while continuing to increase in diameter (phase 2). The simultaneous expansion in length and diameter in phase 1 is possible only until the density of bees approaches a minimum below which the bonds between bees begin to strain and would eventually break. To avoid this limit, the clusters begin to shorten while increasing their diameter in phase 2. This allows them to continue to increase their surface area without a dramatic decrease in density. Alternatively, it is possible that phase 1 of expansion occurs in response to the derivative of temperature with respect to time, suggesting that monotonic increases in diameter and length are a strategy used to respond to increasing temperature rather than the absolute temperature. For the slower 8.6°C/hr and 4.3°C/hr, this transition from phase 1 to phase 2 occurs when the heating ramp ends and the plateau begins, suggesting that phase 2 may be a strategy used in response to high absolute temperatures. If this is true, the transition may be delayed in the fastest treatment (33.4°C/hr) because the internal temperature of the swarm changes more slowly than does the ambient air (Fig. 3D).

The contraction of the swarm in response to cooling also follows a two phase response. At the start of the cooling ramp, the cluster initially decreases rapidly in length and decreases more slowly in diameter (phase I), but when the temperature reaches the 15°C plateau, the cluster begins to increase in length as its diameter continues to decrease leading to a decrease in surface area to volume ratio (phase 2). The rapid shortening of the swarm in phase 1 likely occurs because it is relatively easy for bees to climb up chains of bees toward the center of the swarm to avoid cooling, but it is more difficult for bees at the base to move inward because they are bearing the load of other bees. The transition from phase 1 to phase 2 may occur when the cluster approaches a maximum density beyond which shortening of the cluster is no longer possible. Alternatively, phase 1 may be a response to the derivative of the temperature ramp and phase 2 may be a response to the absolute low temperature during the plateau.

The slow, hysteretic and anisotropic nature of the morphological responses of the swarm clusters to heating and cooling events point to the presence of mechanical constraints on the topological rearrangements of these loaded networks of interconnected bees. The movement of a bee in response to local temperature gradients may be limited not only by the local density of bees but also by the network structure of the cluster and the loading of individual bees. Although the internal structure of swarm clusters has not been rigorously quantified due to obvious methodological challenges, previous observational work has suggested the presence of a dense ‘mantle’ of bees on the surface which push toward the center of the cluster at cool temperatures [9]. The center of the cluster is thought to be composed by vertically oriented chains of loaded bees (which support the swarm) as well as caverns in which other unloaded bees which are free to move around [12, 9].

Peleg et al. [18] recently discovered that swarm clusters are able to change their morphology in response to mechanical perturbations. When a swarm is subjected to horizontal oscillations (similar to those imposed by a tree branch swaying in the wind), the cluster spreads out to increase its attachment area, thereby reducing the overall mechanical strain experienced by the cluster. This collective response requires individual bees to sense local strains and move from regions of low strain to regions of higher strain in order to stabilize the cluster as a whole. Oscillations with low accelerations induce little spreading, while oscillations high accelerations lead to lots of spreading, suggesting a graded response. Taken together, these experiments suggest that morphological responses of swarm clusters to their environmental conditions are multi-functional and require collective multi-sensory integration. Figure 6 illustrates this phase space when the cluster reaches a steady state (*t* = inf). At low temperature and low mechanical forcing the swarm is relatively dense and conical (except when Δ*T*/*T*_0_ *<* 0 [9]). At high temperatures and mechanical forcing the swarm tends to increase its attachment area and flatten out. When temperatures are too high, the cluster may break up (Fig. S1) and when mechanical forcing is too high the cluster may also break up [9].

Taken together, our results suggest that thermoregulatory morphing behavior by honeybee swarm clusters in response to ambient temperature fluctuations is relatively slow, hysteretic, anisotropic and history dependent. As a model system, honeybee swarms can shed light on how stimulus-response behaviors of individuals and physical coupling through a shared environment can give rise to collective physiological processes such as thermoregulation. More empirical work is needed to understand the movements of individual bees, the internal structure of the cluster and the integration of mechanical and temperature cues.

## Author contributions

JMP, OP and LM conceived and designed the study. JMP built the experimental apparatus. JMP and OP conducted the 2017 experiments and JMP conducted the 2018 experiments. JMP and LM analyzed the data and JMP made the figures. JMP wrote the manuscript, and all authors edited it. LM supervised the project. All authors contributed to revisions and gave final approval for publication.

## Data accessibility

Data included in this manuscript are available upon reasonable request.

## Acknowledgements

We thank Jim MacArthur for building the sensor array and valuable advice on instrumentation. We thank Mary Salcedo for her contributions to preliminary experiments that are not included in this write up but were valuable in designing this study.

